# A predicted deleterious allele of the essential meiosis gene *MND1*, present in ~3% of East Asians, does not disrupt reproduction in mice

**DOI:** 10.1101/673145

**Authors:** Tina N. Tran, Julianna Martinez, John C. Schimenti

## Abstract

Infertility is a major health problem affecting ~15% of couples worldwide. Except for cases involving readily-detectable chromosome aberrations, confident identification of a causative genetic defect is problematic. Despite the advent of genome sequencing for diagnostic purposes, the preponderance of segregating genetic variants complicates identification of culprit genetic alleles or mutations. Many algorithms have been developed to predict the effects of “variants of unknown significance” (VUS), typically SNPs (single nucleotide polymorphisms), but these predictions are not sufficiently accurate for clinical action. As part of a project to identify population variants that impact fertility, we have been generating CRISPR-Cas9 edited mouse models of suspect SNPs in genes that are known to be required for fertility in mice. Here, we present data on a non-synonymous (amino acid altering) SNP (rs140107488) in the meiosis gene *Mnd1*, which is predicted bioinformatically to be deleterious to protein function. We report that when modeled in mice, this allele (*MND1*^*K85M*^), which is present allele frequency of ~3% in East Asians, has no discernable effect upon fertility, fecundity, or gametogenesis, although it may cause sex skewing of progeny from homozygous males. In sum, this appears to be a benign allele that can be eliminated or de-prioritized in clinical genomic analyses of infertility patients.

## Introduction

Meiosis begins with programmed formation of double stranded breaks (DSBs) induced by the SPO11 protein (Gray and Cohen, 2016). Processing and repair of these DSBs via recombination is crucial for proper pairing and segregation of homologous chromosomes at the first meiotic division. The recombination repair process involves several steps, including resection of the 5’ ends, loading of proteins (including the RPA complex) that protect the 3’ overhangs, replacement of RPA by the RecA-related recombinases RAD51 and DMC1, then invasion of the single stranded nucleoprotein filament into the same locus of the homologous chromosome to form recombination intermediates called D-loops. Subsequent resolution of the recombination intermediates occurs via non-crossover (NCO) and crossover (CO) recombination. The recombinase loading and D-loop formation depend upon a heterodimer of the MND1 and HOP2 accessory proteins (Bugreev et al., 2014; Chen et al., 2004; Petukhova et al., 2005).

*Mnd1* knockout mice of both sexes are sterile, causing an absence of oocytes and postmeiotic spermatids, respectively (Pezza et al., 2014). Most mutant spermatocytes were defective in homologous chromosome synapsis and DSB repair, leading to apoptosis. However, 23% of spermatocytes exhibited extensive or complete synapsis and underwent HOP2-dependent DSB repair, although they were deficient in crossovers (chiasmata) that are crucial for normal chromosome disjunction (and thus, aneuploidy prevention) (Pezza et al., 2014).

As part of a project to identify SNPs that cause infertility in human populations, we report here on the creation and analysis of a mouse model containing a predicted deleterious non-synonymous (amino acid altering) SNP (rs140107488) in *MND1.* This SNP is present at an allele frequency of 1.6% in East Asians (meaning that ~3.2% are carriers) and 0.14% in South Asians (gnomAD database). We find that mice bearing this SNP are fertile, and show no signs of defective meiosis.

## Materials and Methods

### Mouse generation and genotyping

The DNA template for making sgRNA was generated as described via overlap PCR (Varshney et al., 2015). The guide RNA sequence corresponding to *Mnd1* was: AGCCTCCAACTTGCGCTTCC, embedded within the forward overlap primer: GAAATTAATACGACTCACTATAGG<u>AGCCTCCAACTTGCGCTTCC</u>GTTTTAGAGCTAGAAATAGC. This was matched with a universal reverse primer: CAAAATCTCGATCTTTATCGTTCAATTTTATTCCGATCAGGCAATAGTTGAACTTTTTCACCGTGGCTCAGCCACGAAAA.

The DNA template produced by the overlap PCR was reverse-transcribed into RNA using Ambion MEGAshortscript T7 Transcription Kit (cat#AM1354), and purified using Qiagen MinElute columns (cat#28004). For pronuclear injection, the sgRNA (50 ng/μL), single stranded homologous recombination template (ssODN; 50 ng/μL, IDT Ultramer Service), and Cas9 mRNA (25 ng/μL, TriLink) were co-injected into zygotes (F1 hybrids between strains FVB/NJ and B6(Cg)-Tyr^c-2J^/J) then transferred into the oviducts of pseudopregnant females. The ssODN sequence was as follows: AAAGCACATCTCTTCATAAGGCGGTGACTTACCTGAGAGTTCAGGGCCTCTAACAT GCGCTTCCTTGCATGAAGAGCCTTGCTCGGAAAAGCCCAATAGTAATTGGACGTCCCGA. Founders carrying at least one copy of the desired alteration were identified by PCR using primers flanking the targeting site, followed by Sanger sequencing.

Crude lysates for PCR were made from small tissue biopsies (tail, toe or ear punches) as described (Truett et al., 2000). Genotyping primers were as follows: 5’-GCGTTGAGCCCAAATAAGAA-3’ and 5’-GTGGATGACGGTATGGTTGA-3’. Cycling conditions were: initial denaturation at 95° for 5 min, then 30 cycles of 95° for 30 sec, 58° for 30 sec, 72° for 30 sec, and final elongation at 72° for 5 min. For identification of *Mnd1* mutants, the 233bp amplimer was digested with *Hae*III, and run on an agarose gel. The mutant, but not WT product was cleaved into 2 fragments of 133 and 100bp.

All animal use was conducted under protocol (2004-0038) to J.C.S. and approved by Cornell University’s Institutional Animal Use and Care Committee.

### Immunocytochemistry of spermatocyte chromosomes

This was performed essentially as described, but summarized as follows (McNairn et al., 2017). To prepare surface spreads, testes were isolated from eight-to twelve-week old males, detunicated, and minced in MEM media. Spermatocytes were hypotonically swollen in 4.5% sucrose solution, lysed in 0.1% Triton X-100, 0.02% SDS, and 2% formalin. Slides were blocked for 1 hour at room temperature in 5% goat serum in PBS, 0.1% Tween20. Primary antibodies and dilutions used were: anti-SYCP3 (1:600, Abcam, #ab15093), anti-DMC1 (1:100, Abcam, #ab11054), and anti-MLH1 (1:100, BD Pharmingen, #554073). Immunolabeling was done overnight at 4ºC. Secondary antibodies used were: goat anti-mouse IgG AlexaFluor 488 (1:1000, ThermoFisher Scientific, #R37120) and goat anti-rabbit IgG AlexaFluor 594 (1:800, ThermoFisher Scientific, #R37117). Secondary antibodies were incubated for 1 hour at room temperature. Foci were quantified using ImageJ with plugins Cell Counter (Kurt De Vos) and Nucleus Counter. All data was analyzed in Prism 7 (GraphPad).

### Sperm counts and histology

For sperm counts, cauda epididymides (both sides) were collected from 8-week old males, minced in PBS, then incubated at 37ºC for 10 minutes to allow sperm to swim out. Sperm solutions were diluted and counted on a hemocytometer.

For histological sections, testes were fixed in Bouin’s for 24h, washed in 70% ethanol for 24h, then embedded in paraffin. Sections were made at 6μM thickness. Slides were stained with hematoxylin and eosin (H&E).

## Results and Discussion

### Selection of the *MND1*^*K85M*^, encoded by SNP variant rs140107488, as a potential infertility allele

In our ongoing project to identify and functionally validate segregating infertility alleles in human populations by mouse modeling, we select candidate non-synonymous SNPs (nsSNP) in genes that are known to be required for fertility in mice based on knockout studies, and which are computationally predicted to be pathogenic. rs140107488 encodes a lysine-to-methionine change at amino acid (AA) position 85 of human and mouse MND1. The SIFT, PolyPhen-2, and Mutation Assessor algorithms, which predict deleteriousness of an nsSNP based on physico-chemical properties of the substituted AA, potential structural impact, and evolutionary conservation (Adzhubei et al., 2013; Kumar et al., 2009; Reva et al., 2007), all predict this variant to be highly deleterious to protein function. These algorithms’ scores for rs140107488 are, respectively, 0 (the “worst” possible), 0.963 (“probably damaging”) and 0.933 (“high” likelihood of being damaging). Given these predictions, in conjunction with the knowledge that MND1 is essential for meiosis and exists as a significant minor allele in East Asians, we selected it to model in mice and assess the consequences *in vivo*.

### *Mnd1*^*K85M/K85M*^ mice are fertile and produce normal numbers of sperm

To generate mice modeling the human *MND1*^*K85M*^ allele, CRISPR/Cas9-mediated genome editing in single-celled zygotes was performed. An ssODN was used as a repair template to introduce a nucleotide change into codon 85 via homologous recombination, causing this codon to encode methionine instead of lysine (Fig. 1a). Other silent changes were also introduced to facilitate genotyping (creation of a novel restriction enzyme site) and to reduce deletion mutations (mutation of the PAM site required for Cas9:sgRNA recognition) (Singh et al., 2015). Founder mice with the correct mutation were identified (Fig. 1b), backcrossed into FVB/NJ for at least two generations, then intercrossed to generate homozygous mice for phenotypic analysis.

**Figure 1.**
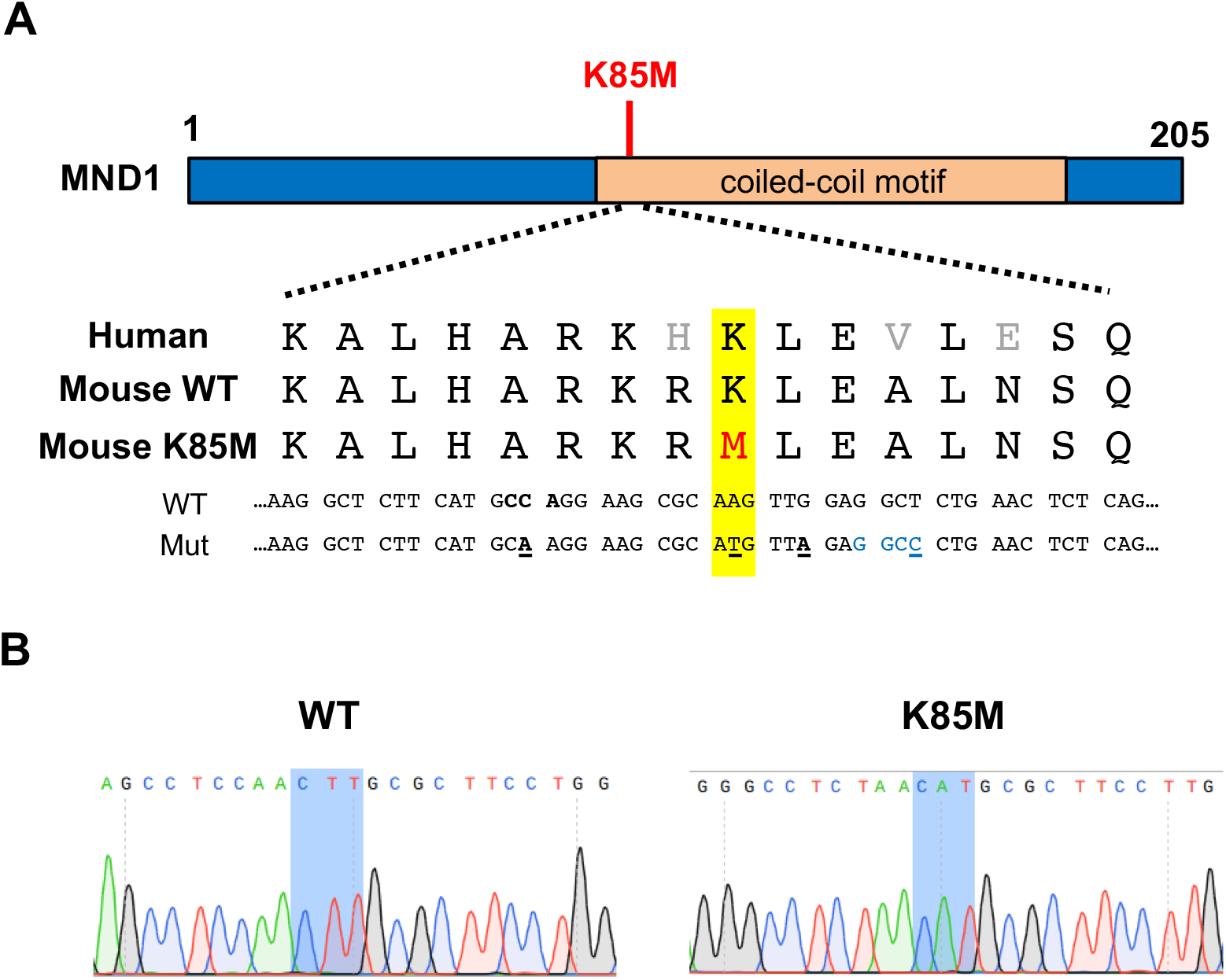
CRISPR/Cas9 genome editing strategy to generate mice *Mnd1*^*K85M*^. A) Schematic of the human MND1 protein and amino acid and nucleotide sequences surrounding the target SNP in the human and mouse genomes. Point changes introduced by the ssODN/HR template are indicated as follows: yellow highlight, red text/underline = nucleotide change(s) to generate the SNP variant; black text/underline = silent nucleotide changes to prevent Cas9 re-cleavage, including the PAM site (also bolded); blue text = *Hae*III restriction enzyme site used for genotyping. K85M, MND1^K85M^. B) Chromatograms of WT and *Mnd1*^*K85M/K85M*^ mice (sequence is reverse complement of the sequences above).

Homozygous mutant mice were normal in gross appearance, consistent with the lack of somatic phenotypes reported for null mice (Pezza et al., 2014). To determine if the *Mnd1*^*K85M*^ allele impacts fertility, homozygotes and wild-type (WT; +/+ or heterozygotes from the same genetic background) adults between 2-7 months of age were bred to WT mates, and litter sizes were recorded. The average litter sizes produced by *Mnd1*^*K85M/K85M*^ males and females were 8.2 (n=123 pups) and 8.6 (n=95 pups), respectively, compared to 8.3 for WT intercrosses (n=58 pups). Thus, fecundity of mutants was normal (p = 0.78 and 0.93 for females and males, respectively by T-test). Interestingly, however, male (but not female) homozygotes produced more male (82) than female (41) offspring (p=0.000073 by T-test), a 2:1 ratio (Fig. 1a). The basis for this is unknown.

To determine if there were potential subclinical effects in sperm production or spermatogenesis in mutants, we performed gross and histological analyses of testes and sperm. Eight-week old *Mnd1*^*K85M/K85M*^ males had testis weights and epididymal sperm numbers that were similar to WT (Fig. 2b). Consistent with these observations, testis histological cross-sections appeared normal in the mutant; all spermatogenic stages were present, and no notable pathologies or abnormal cellular phenotypes were observed (Fig. 2c).

**Figure 2.**
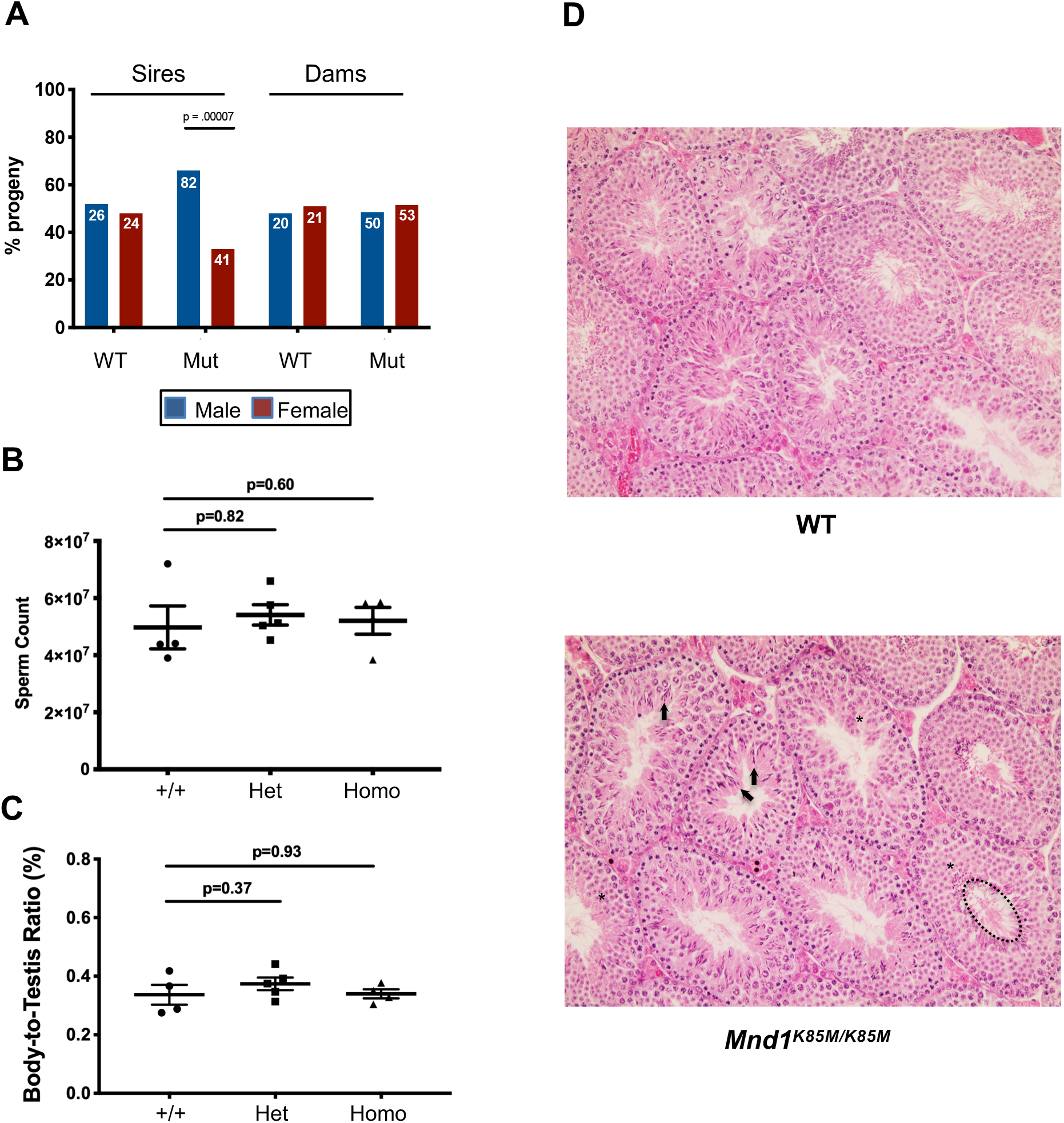
*Mnd1*^*K85M/K85M*^ males have normal sperm counts and spermatogenesis, but exhibit sex distortion. A) Progeny separated by sex of WT and *Mnd1*^*K85M/K85M*^ mice mated to WT animals. Note that mutant males (sires) produce a 2:1 ratio of male:female offspring. B) Total sperm quantified from both epididymides of each animal. N=3-5 per genotype. C) Relative body:testis masses. N=4-5 per genotype. D) Testis histology. H&E stained testis cross-sections of 8-week old males. Highlighted in selected seminiferous tubules of the mutant sample are normal examples of elongated spermatids (arrows), round spermatids (asterisks), and flagellated, spermiated spermatozoa (dashed ellipse in lower right tubule).

MND1 is an important binding partner of HOP2, and this complex stabilizes DMC1 and RAD51 binding to ssDNA to promote strand invasion into, and recombination with, the homologous chromosome (Tourtellotte et al., 1999). To assess if MND1^K85M^ affects DMC1 recruitment and displacement, spermatocyte chromosome surface spreads were immunolabeled with DMC1 and the synaptonemal complex (SC) axial element protein SYCP3 (Fig. 3a). In response to meiotically programmed DSBs, DMC1 loading were observed in foci that were present at normal levels (compared to WT; ~200 foci/nucleus) in leptotene spermatocytes (cells in which homologous chromosome pairing and synapsis has yet to occur). Focus numbers also disappeared normally as meiotic prophase I progressed into pachynema (when chromosome synapsis is complete), indicating that homologous recombination-mediated DSB repair was occurring without defects in the mutant (Fig. 3b). Unlike *Mnd1* null spermatocytes, we did not observe defects in chromosome synapsis (Pezza et al., 2014). To assess crossing-over, late pachytene spreads were immunolabeled with the chiasmata marker MLH1. In contrast to null spermatocytes that have decreased crossing-over in surviving spermatocytes (Pezza et al., 2014), there was no significant difference in MLH1 focus numbers between WT and *Mnd1*^*K85M/K85M*^ (p=0.13) (Fig. 3c, d), indicating a normal level of crossing over. Aberrations in crossover number can lead to aneuploidy and death of offspring, typically resulting in reduced litter sizes, which is not the case with *Mnd1*^*K85M*^ mutants, as indicated above.

**Figure 3.**
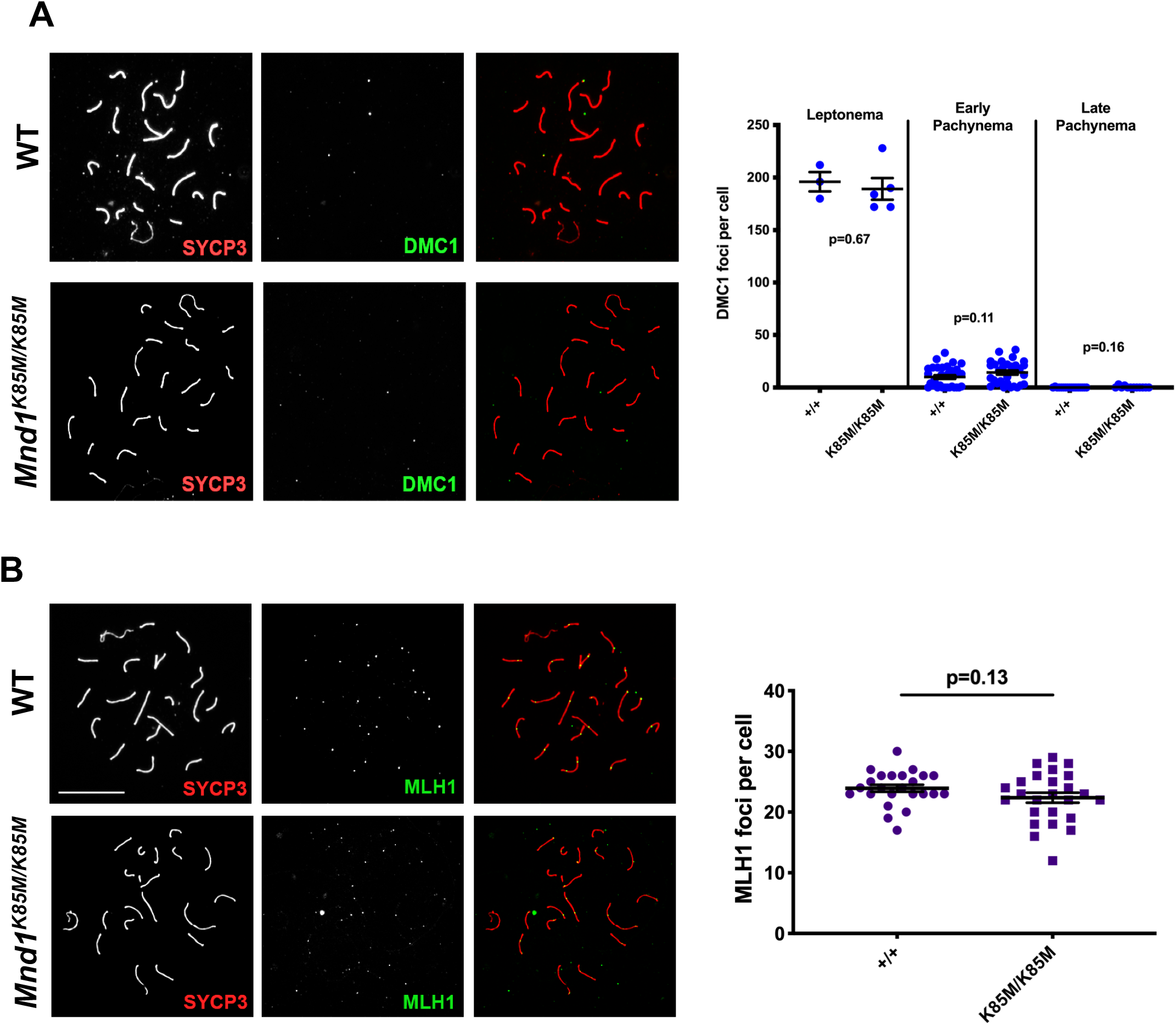
Cytological data indicating that *Mnd1*^*K85M/K85M*^ is not defective in meiotic recombination. A) Late pachytene spermatocyte chromosome surface spreads immunolabeled with SYCP3 and DMC1 (merged in rightmost panels). Numbers of DMC1 foci at indicated substages are plotted. N=2 per genotype. P values are from student’s T-test. Size bar = 20 μm. B) Spreads immunolabeled with MLH1 and SYCP3 (merged in rightmost panels). Size bar = 20 μm. D) Quantification of MLH1 foci in pachytene cells. N=3 per genotype.

In summary, our studies indicate that *MND1*^*K85M*^ is a benign allele that has no discernable effect upon fertility. Homozygous mutants of both sexes were normal in both reproductive and molecular phenotypes. By eliminating this as a candidate infertility allele, these finding may be useful in prioritizing candidate genetic variants that may underlie reproductive infertility in people bearing this SNP. While it is conceivable that the allele is pathogenic in humans but not mice, the consequences are likely to be subtle or subclinical, especially since the allele is present at a relatively high frequency in East Asians. One remaining question is the unusual sex distortion ratio exhibited by allele transmission from homozygous males. At present, it is unclear whether this is a consequence of the mutant allele itself, a statistical rarity, or perhaps an unrelated closely linked allele. Further studies of this phenomenon are under way.

## Acknowledgements

We thank R. Munroe and C. Abratte of Cornell’s transgenic facility for generating the edited mice.

## Authors’ roles

T.T. conducted most of the experiments, wrote a draft of the text and figures, and supervised JM, who contributed to the genotyping and immunhistochemistry analyses. JS supervised the work and the aforementioned students, and prepared the manuscript.

